# Spatiotemporal, environmental, and behavioral predictors of *Varroa* mite intensity in managed honey bee apiaries

**DOI:** 10.1101/2024.09.23.614412

**Authors:** Laura F. Boehm Vock, Lauren M. Mossman, Zoi Rapti, Adam G. Dolezal, Sara M. Clifton

**Affiliations:** Department of Mathematics, Statistics, and Computer Science, St. Olaf College, Northfield, Minnesota, USA; Department of Mathematics, University of California Davis, Davis, California, USA; Department of Mathematics, University of Illinois at Urbana-Champaign, Urbana, Illinois, USA; Department of Entomology, University of Illinois at Urbana-Champaign, Urbana, Illinois, USA; Department of Mathematics, Denison University, Granville, Ohio, USA

## Abstract

Honey bees contribute substantially to the world economy through pollination services and honey production. In the U.S. alone, honey bee pollination is estimated to contribute at least $11 billion annually, primarily through the pollination of specialty crops. However, beekeepers lose about half of their hives every season due to disease, insecticides, and other environmental factors. Here, we explore and validate a spatiotemporal statistical model of *Varroa destructor* mite burden (in mites/300 bees) in managed honey bee colonies, exploring the impact of both environmental factors and beekeeper behaviors. We examine risk factors for *Varroa* infestation using apiary inspection data collected across the state of Illinois over 2018-19, and we test the models using inspection data from 2020-21. After accounting for spatial and temporal trends, we find that environmental factors (e.g., floral quality, insecticide load) are not predictive of *Varroa* intensity, while several beekeeper behaviors (e.g., smaller colony density, supplemental feeding, and mite monitoring/treatment) are protective against *Varroa*. Interestingly, while monitoring *and* treating for *Varroa* is protective, treating *without* monitoring is no more effective than not treating at all. This is an important result supporting Integrated Pest Management (IPM) approaches.

**Author Summary:** Honey bees contribute substantially to the world economy through pollination services and honey production. However, beekeepers lose about half of their hives every season due to disease, insecticides, and other environmental factors. Pathogens, such as *Varroa* mites and the viruses they vector, are especially detrimental to colony health, and best practices for pest management remain contentious. In this study, we model *Varroa destructor* mite burden in managed honey bee colonies using apiary inspection data collected across the state of Illinois from 2018 – 2021. Our modelling approach accounts for both spatial and temporal trends, allowing us to investigate the marginal impacts of environmental factors and beekeeper interventions on mite burden. We show that treating for *Varroa* mites has a protective effect only when accompanied by a monitoring strategy, important evidence in favor of Integrated Pest Management (IPM) approaches.

## Introduction

Pollinators face substantial environmental challenges, often classified into four main stressors: parasites, pathogens, pesticides, and poor nutrition (the four P’s) [1]. Poor environmental conditions lead to weakened or even collapsed colonies, which has serious ecological and economic consequences [2–4]. An estimated 87.5% of flowering plants are pollinated by insects and other animals [5], and pollination services are estimated to contribute at least $235 billion annually in the world economy [6–8]. Honey bees (*Apis mellifera*) are of particular economic interest because the species is one of only a few pollinators domesticated for honey production and crop pollination [9].

Because of the importance of pollinators to the environment and economy, colony failure has attracted considerable attention from mathematical and statistical modelers. While the role of parasites and pathogens, such as *Varroa* mites and the viruses they vector (e.g., Deformed Wing Virus and Acute Bee Paralysis Virus) [10–12], remain the most studied culprits, other factors, such as pesticides [13] and poor nutrition [14], have also been considered. Every modeling effort has simplified the inherently complex pollinator system by focusing on a few aspects of the disease system while de-emphasizing other aspects.

For example, Kang et al. model bee-to-bee virus transmission explicitly by dividing the populations of both bees and mites into susceptible and infected [10]; seasonal effects, the role of nutrition, and mite dispersion among colonies, are not considered. Other models focus attention on colony demography, seasonality, and food resources, while ignoring disease spread [14–16]. Still other models address disease and demographic dynamics, while ignoring environmental conditions [12, 17, 18].

While within-hive disease dynamics have been modeled with various levels of complexity [19], between-hive transmission over large spatial areas has been mostly neglected. This is understandable because large-scale colony parasite monitoring by single agencies, collecting and sharing data in a consistent manner, is sparse in space, time, and/or covariates. For example, the Bee Informed Partnership has performed the most widespread honey bee colony surveys over the last decade [20, 21]; however, these surveys focus primarily on annual and seasonal colony losses and do not usually incorporate more complex factors, practices, or exact location data. Other recent studies have modeled the transmission of various parasites and pathogens within a single apiary [22, 23], but to our knowledge very few modeling efforts (e.g., [24]) examine risk factors for parasites and pathogens (e.g., environmental conditions and beekeeper behaviors) over large spatial areas.

To fill the void on large spatial scale risk factors for *Varroa* infestation in managed honey bee colonies, we build and validate several spatiotemporal statistical models of *Varroa* intensity (mites/300 bees) in colonies across the state of Illinois. To train and test the models, we merge several large-scale datasets of apiary inspections, beekeeper surveys, and environmental factors. After accounting for spatial and temporal trends (our baseline model), we compare the marginal effects of many factors hypothesized to increase risk for or protect against mite infestation. These factors include environmental conditions, such as floral quality, nesting quality, insecticide load, and apiary density.

We also explore the impact of beekeeper interventions, including supplemental feeding, parasite monitoring, and various *Varroa* treatments.

## Methods

### Data

Our spatiotemporal data fall into three general categories: colony health status (with a location and a date), beekeeper behaviors (with a location and a year), and environmental indicators (with a location, sampled in 2018 but assumed to be stable over several years).

These data are collected from two main sources: (1) colony health status and beekeeper behaviors are available via yearly inspections in the state of Illinois from 2018 to 2021 [25], and (2) environmental indicators are available from Beescape (beescape.psu.edu). These sources cover all four of the main pollinator stressors (parasites, pathogens, pesticides, and poor nutrition) [1]. Because parasites and pathogens are highly correlated in our sample (see Figure 1), we will use *Varroa* intensity as a proxy for both.

**Fig 1.**
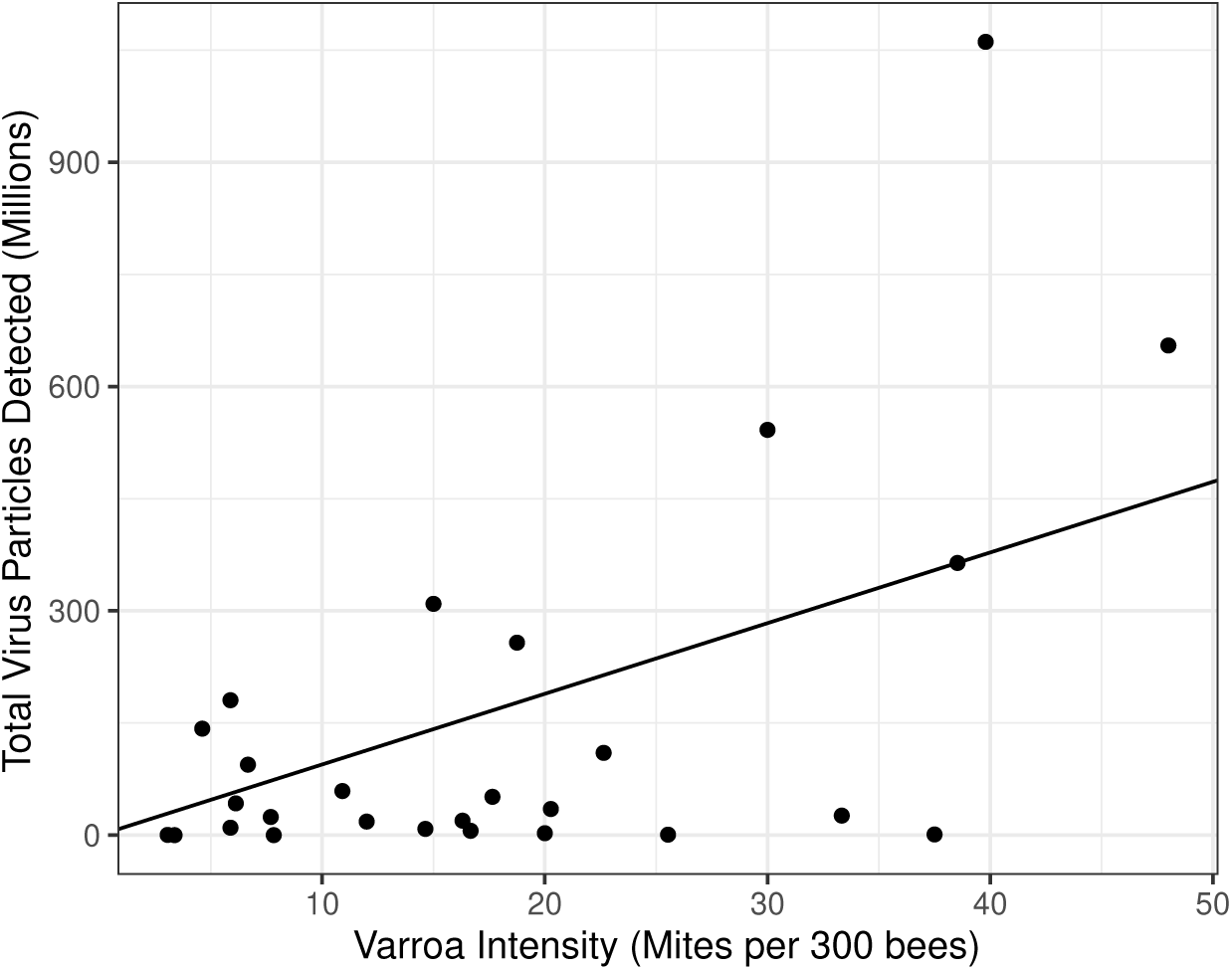
Mite burden is correlated with viral burden (model *R*^2^ = 0.50) [25].

The Illinois inspection data are thorough; each hive inspection includes the spatial location, multiple hive health indicators (e.g., queen status, amount of honey, egg/larvae presence), infestation status (e.g., *Varroa* mites per 300 bees, American or European foulbrood detection, small hive beetle presence), and disease control indicators (e.g., prophylactic treatments, beekeeper monitoring frequency and method, and intervention frequency and method).

Although the parasite and pathogen data are thorough, they present several challenges and uncertainties. First, the inspection data are sparse in time; inspections of each hive occur only once per year, and only about one in five registered apiaries are inspected (it should be noted that this is a huge fraction of apiaries inspected relative to similar efforts, e.g., [20, 21]). Second, while registration and inspections are mandated in Illinois, a non-negligible number of data points are missing important features (e.g., a valid location, date, or disease status); see Data Cleaning section for details. Third, *Varroa* in particular must infest at least 1% of the hive to be detected; the standard method for testing is to remove and inspect 300 adult bees from a hive, and the limit of detection is one mite per sample with an assumed correction factor of 2-3 times for pupae (as pupae can be parasitized but cannot be sampled via this method) [26]. Finally, each data point is generated by a human inspector, and therefore may have errors or uncertainties that are not easily quantified.

Data relevant to pesticides and poor nutrition were scraped from the Beescape Map Tool [27] at each reported hive location (latitude and longitude) from 2018 inspections [28]. This application scores forage (floral) quality in spring, summer, and fall; nesting quality; and insecticide load [27] using a rigorously validated model for wild bee abundance [29, 30].

The forage quality score in each season is a weighted average of the density and supply of floral resources within a 3 km foraging radius of each hive location. The index, ranging from 0 to 100, is based on satellite imaging of natural areas and USDA crop surveys. Because each colony is inspected only once per year, sometime between May and October, and because all three seasonal floral indicators (*f_spr_, f_sum_, f_fall_*) are highly correlated (from 0.91 to 0.99, depending on the pair) we use the estimated average floral quality over that six month period via the trapezoid rule assuming spring to summer and summer to fall are three months each (see Figure 2 for a visualization):

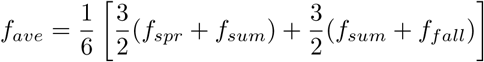

**Fig 2.**
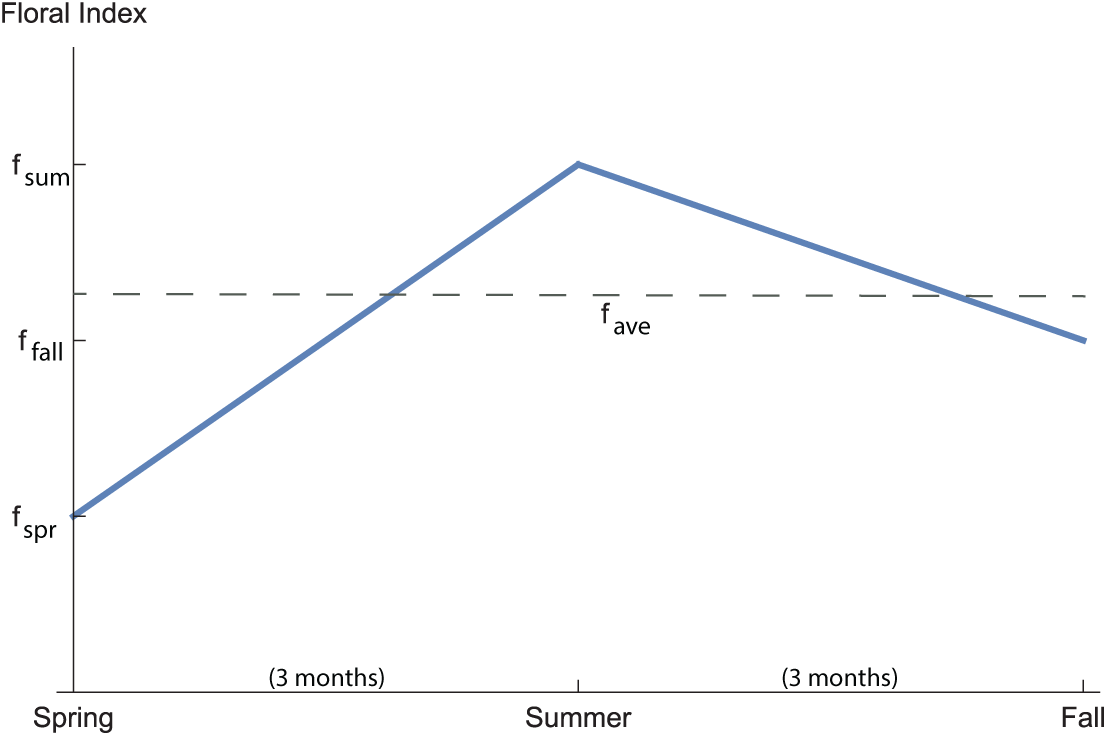
We use a floral indicator which is an estimated average of the seasonal floral indicators over the inspection period May through October. To estimate the average floral quality using the trapezoid rule, we assume spring to summer and summer to fall are three months each.

The nesting quality is estimated by averaging expert opinion on wild bee nesting quality provided by each land cover type. The nesting index, also ranging from 0 to 100, is not expected to be relevant for managed bees with man-made hives, and therefore serves as a check on our results.

The insecticide load score captures the expected toxic load of insecticides applied surrounding each reported hive location. The insecticide score is a weighted average of the lethal doses per area, scaled to fall in the same range as the nesting and forage indices. Theoretically, the insecticide score could take on any non-negative value, but the interquartile range is 79 to 226 across four representative U.S. states (Pennsylvania, Indiana, Illinois, and West Virginia).

While we would have preferred to scrape the Beescape Map Tool each year to establish time-stamped environmental indicators at each inspected hive, the API became inaccessible after changing hosts around 2020. Therefore, we use the scraped environmental data from hive locations inspected in 2018 [25], and we spatially interpolate the indicators for locations inspected in subsequent years, assuming that the indicators remain relatively stable (see Figure 3).

**Fig 3.**
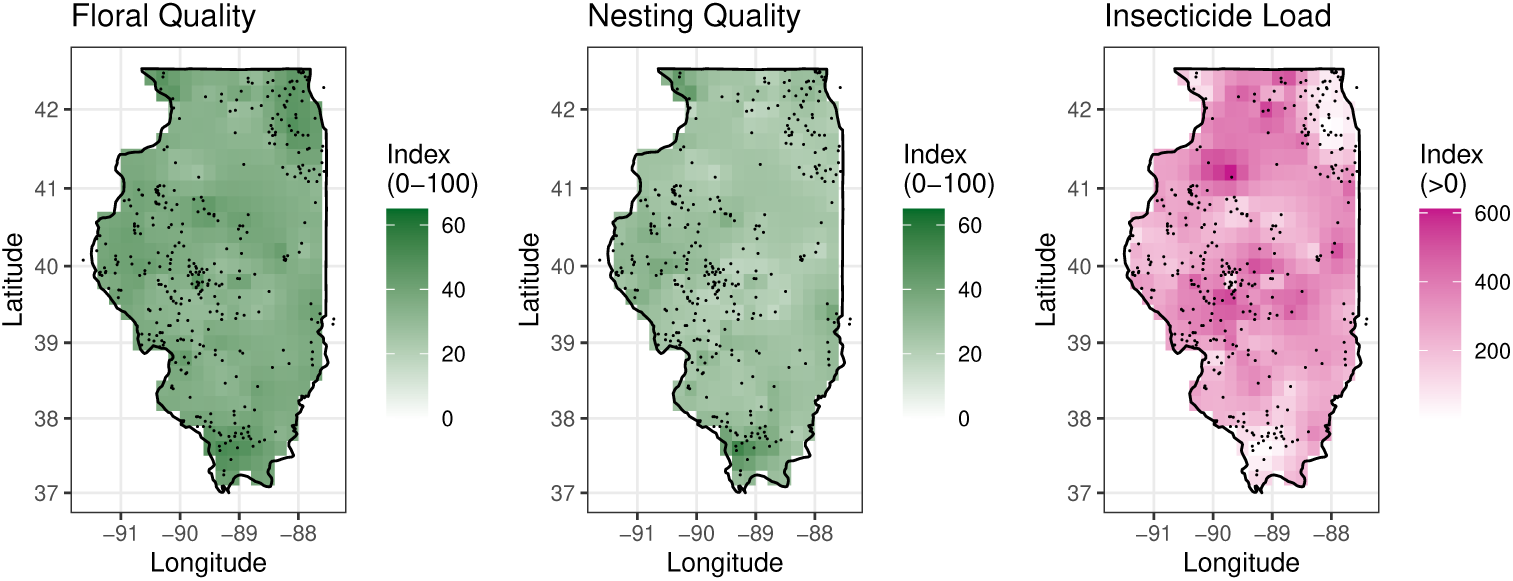
Environmental indicators, interpolated using kriging [31]. Black dots are colony locations in 2018 - 2021.

### Data cleaning

The inspection data contain 1585 recorded inspections in the years 2018-2021. We removed all inspections without dates, which reduced the number to 1546. Of these, 1251 have valid latitude and longitude recorded. For an additional 132 observations, we pull the latitude and longitude from the 2018 Colony Report by matching the registration ID. These values indicate individual hive information. We combine observations at multiple hives from the same apiary (based on shared Latitude/Longitude) into a single value per apiary. If the *Varroa* intensity is measured at more than one hive at a single apiary, we calculate the average intensity value.

In total, this results in 269 unique apiaries in 2018–2019 which we use as training data for the models, and 140 apiaries in 2020–2021 which are used for model testing (see Tables 1 and 2 for summary statistics).

**Table 1.** Summary statistics for *Varroa* intensity in number of mites per 300 bees and potential environmental explanatory variables, Mean (SD). Note that the standard deviation for floral quality and insecticide load are lower for testing data than training data due to location variability. Training data (2018-19) apiaries are located throughout the entire state. In 2020, there are hives near the Chicago area and in Central-Western IL only, and in 2021 only near Chicago. The testing data (2020-21) do not include any apiaries in northwest or southern IL. Floral quality and insecticide load are more variable in southern IL in particular (see Fig 3), decreasing the SD for the testing data.

| Variable | 2018-19<br>train | 2020-21<br>test |
| --- | --- | --- |
| Total apiaries | N = 269 | N = 140 |
| Num. mites / 300 bees | 4.7 (5.9) | 4.9 (6.0) |
| Floral Quality | 38 (6.5) | 38 (3.1) |
| Insecticide Load | 255 (160) | 256 (65) |
| Number of Colonies within 5 km | 8 (9) | 7 (9) |

### Baseline model

As *Varroa* is known to increase throughout the summer season [24, 32–34], we first consider a baseline model that incorporates this seasonal effect [35]. Additionally, our initial data exploration suggests that *Varroa* intensity also varies spatially. A Generalized Additive Model (GAM) allows us to flexibly model the nonlinear relationship between intensity and location [36]. Because intensity is measured as a count per 300 bees, we assume the response variable follows a Poisson distribution.

As not all intensities are reported as integers (due to both the averaging of multiple colony values per apiary, and some values being reported in fractions) and to account for the over-dispersion present in the data, we use a negative binomial model. Like the related Poisson regression model, a negative binomial model is suitable for count or rate data, but relaxes the Poisson assumption that the mean is equal to variance [37]. Instead, the variance of the distribution is related to the mean *µ* and the over-dispersion parameter *k*, such that *Var*(*Y*) = *µ* + *µ*^2^/*k*.

The negative binomial regression model is written as

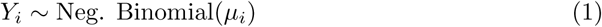

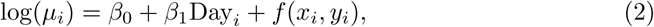

where *Y_i_*is the *Varroa* intensity (averaged over all hives in the apiary) in mites/300 bees at each apiary *i*, *µ_i_* is the mean (or expected) *Varroa* intensity, Day is the number of days since the beginning of the year (January 1 is Day 1) of the apiary inspection, and (*x_i_*, *y_i_*) are the longitude and latitude coordinates of the apiary. The relationship between Day and *log*(*Y_i_*) is assumed linear because exponential growth of *Varroa* intensity has been observed in previous studies [32], and a spline model is not significantly better than the linear model (*p* = 0.964). The function *f* is a low-rank thin plate regression spline, chosen because it has certain optimality properties for multidimensional (e.g., latitude and longitude) smoothing [38].

In our baseline model, we assume errors are not correlated spatially or temporally at a scale we can detect with our sparse data. After accounting for location and time of year, residuals are not spatially or temporally correlated, except possibly at very short spatial scales (< 0.5 km).

All models were fit using the mgcv package in R [39, 40] with smoothing parameters chosen via cross-validation.

### Stepwise regression

We first investigate the effect of each individual beekeeper and environmental variable, after accounting for location and time of year, by adding variables one at a time to the baseline model. We investigate both additive models, in which the effect of the variable is assumed to result in an increase in *Varroa* which is constant across time, as well as an interaction of each variable with day of year, which would result in different rates of growth of *Varroa* through the year.

Next, we investigate variables which were found statistically significant individually by adding them one at a time into a multiple linear regression model. As there were a small number (only treatment, supplemental feeding, and colony density) we consider all three possible pairings and a fourth model which included all three variables.

We evaluate model fit with *R*^2^, AIC, Relative error, and Mean absolute error on the training data (2018-2019) and model prediction with Relative error and Mean absolute error on the test data (2020-2021). We further include the p-value comparing the listed model to the baseline model using a likelihood ratio test for the training data.

Relative error is

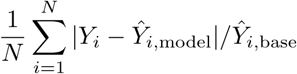

Mean absolute error is

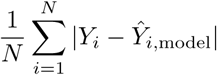

where 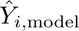 and 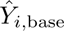 indicated the predicted *Varroa* intensity from the specified model and the baseline model.

AIC is the Akaike Information Criterion, which measures prediction error and is a relative measure of model quality which adjusts for the number of model parameters. The criterion is measured on a log scale, with lower values indicating better fit. Differences of >10 are considered large [41], while differences of <2 are considered negligible.

## Results

### Baseline model

In Figure 4 we see the predictions from the baseline model in each month over the entire state of Illinois. The predicted surface is fit with the 2018-19 training data on the first day of each month. We can compare the observed *Varroa* intensities in the training data (2018-19) and in the test data (2020-2021) to the surface predicted by our model. To see the smooth trend in time, we predict the *Varroa* intensity at the center of the state (−89.68° longitude, 39.90° latitude) each day from May 1 to October 31 (Figure 5).

**Fig 4.**
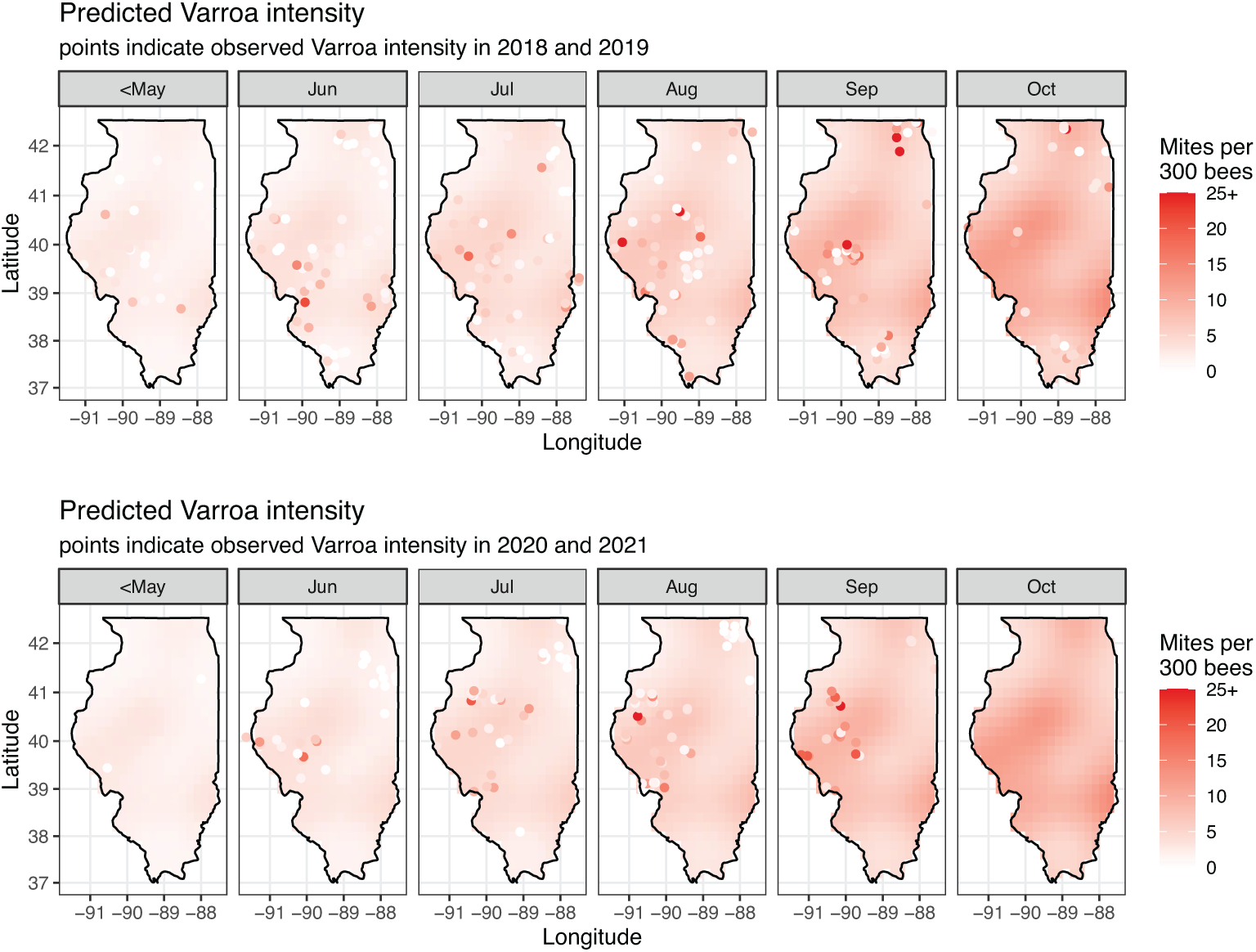
*Varroa* intensity predicted from baseline model (i.e., the model using only day of year and spatial location) using 2018-2019 data. Predictions are for the first day of indicated month. Top row: predicted intensity with observed 2018-19 intensities overlaid (training). Bottom row: predicted intensity with observed 2020-2021 intensities overlaid (testing).

**Fig 5.**
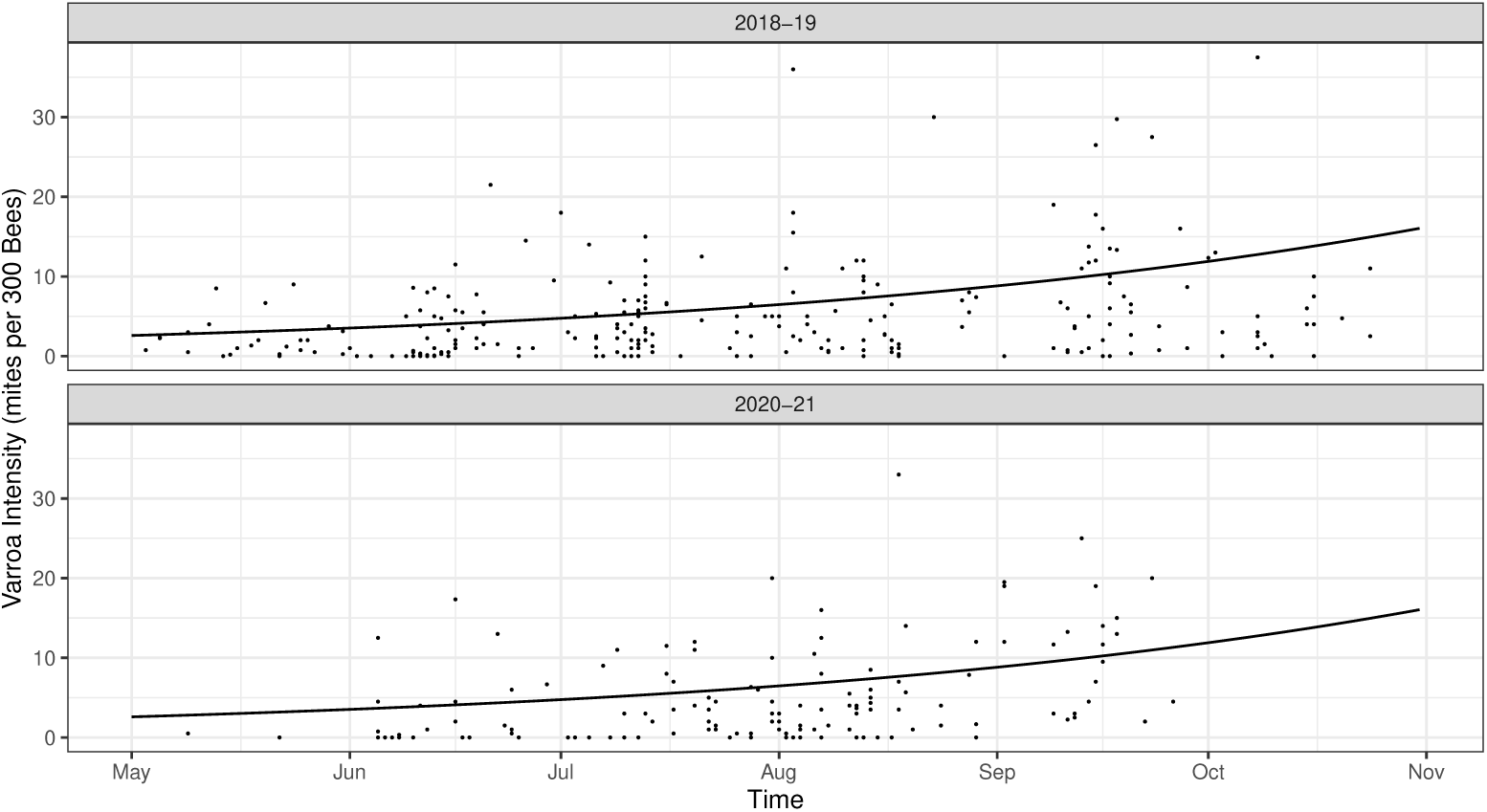
*Varroa* intensity predicted from baseline model at the center of the state of Illinois (−89.68°, 39.90°) using 2018-2019 data (solid line). Dots indicate observed *Varroa* intensities. Top row: predicted intensity with observed 2018-19 intensities overlaid (training). Bottom row: predicted intensity with observed 2020-2021 intensities overlaid (testing).

### Impact of environmental factors

The environmental variables tested are floral quality, insecticide burden, nesting quality, and colony density.

Floral quality, insecticide use and nesting quality are not significant as shown in Table 3. While the additive model for colony density, measured as the number of colonies within a 5 km radius, does not show statistical significance, the model which includes and interaction between day of year and colony density is a significant improvement over the baseline model. This suggests that the growth rate for *Varroa* increases as colony density increases; however the estimated initial *Varroa* intensity at May 1st is lower in regions with higher colony density. Note that in our sample, there does not seem to be an association between treatment strategy and colony density, either globally or regionally. The predicted level of *Varroa* is lower at high density regions until mid August, and then is higher than in low density regions. Graphical results indicate this relationship could be driven by influential late season observations in very high density regions (see Supplemental Materials). The model with density and its interaction with time has the lowest relative error and mean absolute error, both for the training and testing data, indicating that colony density is important for *Varroa* intensity prediction.

### Impact of beekeeper interventions

The majority of the managed apiaries in our dataset (214, or 80%) provide supplemental feeding. Of these, the most common supplement is syrup only (138), or syrup in combination with pollen (29), solid (17), or both pollen and solid (15). Only 15 total apiaries provided supplemental feeding of solids, pollen, or combination, without syrup.

The rate of growth of *Varroa* intensity is 2.4% per day in apiaries that do not supplement, and only 0.8% per day in apiaries that supplement (p-value for test of difference is 0.0007).

The practice of supplemental feeding does not seem to vary across the state; the predominant practice (80% of apiaries observed in 2018-2021) across all regions is to provide supplemental feeding; this makes sense as feeding is widely recommended at some points of the year and is therefore a typical beekeeping practice [42–44]. The model with supplemental feeding has the lowest AIC, indicating good fit to the training data, and the low p-value indicates the observed relationship is statistically significant. It does not however improve model prediction as well as colony density, perhaps because supplemental feeding is so nearly uniformly practiced.

Among many possible *Varroa* management strategies, we consider three categories: “Monitor + Treat”, “Treat only” and “None”, with the monitor and treat strategy being predominant, as seen in Table 2. Due to small sample size, we did not include a “Monitor only” category. There were only 10 apiaries which reported monitoring, but not treatment in 2018-2019. Of these, three had responses to other variables which indicated they may possibly treat (e.g. a last treatment date of 2017, reported treatment frequency of >0 in last 12 months). Therefore these 10 apiaries are included in the “Monitor + Treat” category. As a sensitivity analysis, we repeated the analysis with these 10 apiaries in the “None” (no treatment) category, and achieved similar results. Integrated Pest Management (IPM) approaches recommend only treating in conjunction with testing, in order to reduce ecological contamination and pesticide resistance [45]; therefore, any beekeepers treating without testing are not following IPM best practices.

**Table 2.**
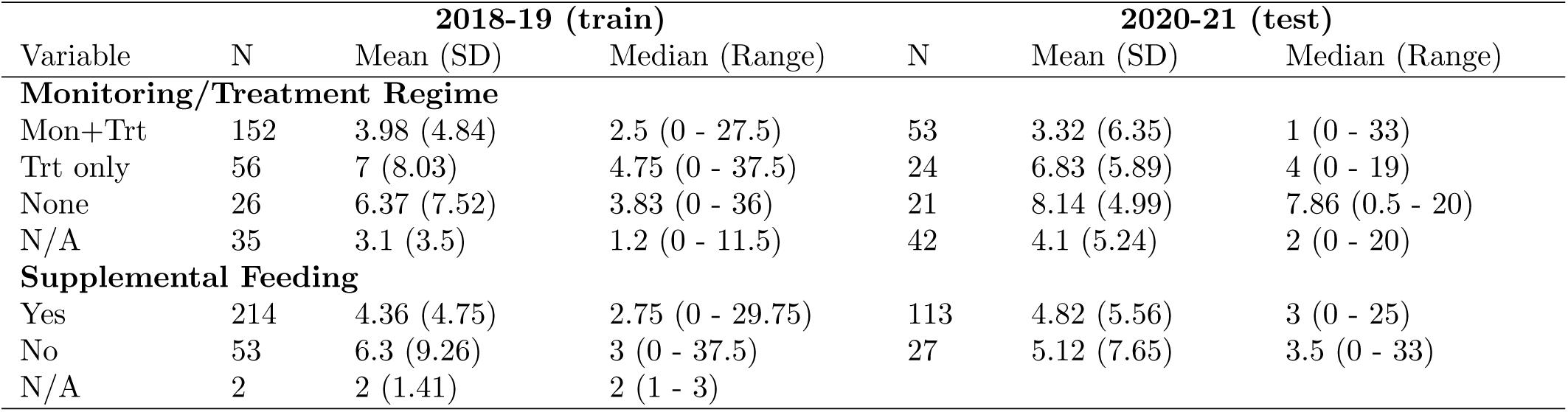
Number of apiaries and summary statistics for *Varroa* intensity (number of mites per 300 bees) by monitoring/treatment regime and by supplemental feeding regime. N/A indicates response is missing in inspection report.

The interaction model shows that the growth rate of *Varroa* over time is significantly higher for the “Treatment only” group compared to the “Monitor + Treatment” strategy (p value = 0.0032). The growth rate for “Monitor + Treat” is 0.5% per day, but for “Treatment only” is 1.7% per day. The “No treatment” group is not significantly different from either of the other groups, although this may be at least partially due to small sample size in that group. Because the sample size on the “no treatment” group is small, the uncertainty in the estimated effect is quite large (it has large standard error). The growth rate for the “no treatment” group is also estimated in between the other two groups (about 1% per day). Since these differences are smaller, and the standard error is larger, the “no treatment” group is not significantly different from either Trt only or Monitor+Trt.

When we test the overall significance of the interaction versus the baseline model, the result appears not significant (p-value = 0.824, Table 3). This somewhat misleading however, as the spatial component of the model is reduced in complexity once we account for *Varroa* management strategies, which vary considerably by location. For example, in central Illinois, where predicted *Varroa* intensity is high, there are a variety of monitoring and treating practices. In Southern Illinois, where predicted intensity is low, nearly everyone monitors.

**Table 3.**
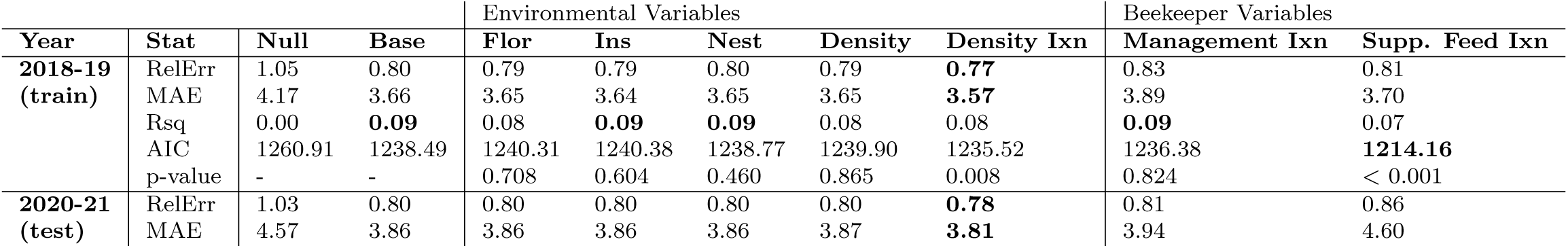
Results of single variable regression models. Relative Error (RelErr = 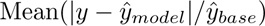, Mean Absolute Error 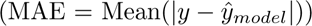. The null model uses overall mean Varroa intensity as the predicted value 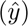. The “Base” model uses only space and time. Columns labeled Ixn include an interaction between the listed variable and time. We also tested interaction models with Floral, Insecticide and Nesting but they were also not statistically significant.

When we compare AIC, we see that the AIC is only slightly lower for the interaction model (1236) compared to baseline (1238), however the net change in model complexity is in favor of the interaction model; though we add terms to the linear part of the model, the reduction in complexity of the spatial fit leads to a model with lower total degrees of freedom. Therefore, we would conclude in favor of the interaction model.

### Multiple regression model

Based on our stepwise regression results, we have identified three variables which are individually important for predicting *Varroa* intensity: apiary density, *Varroa* management strategy, and supplemental feeding practice. We test each combination of these variables to find the best model. Because of the noted relationship of management practice and the spatial component, we compare the 7 potential models with (Table 4) and without (Table 5) the spatial component. There is no clear winner amongst the models for all criterion. We once again observe that the impacts of management strategy are confounded by spatial location. When the spatial component of the model is included (Table 4), including management strategy does not lead to improvements in model fit, regardless of whether colony density or supplemental feeding are accounted for. However, if the spatial component is excluded (Table 5), management strategy is important, particularly when considered in conjunction with supplemental feeding. Because of the spatial patterns in management strategy mentioned in the previous section, it is difficult to disentangle these effects. We do see however, that colony density is important after accounting for this spatial heterogeneity either through inclusion of the spatial component (Table 4) or management strategies (Table 5), both in improving the model fit for the 2018-19 data, and in predicting *Varroa* intensity in 2020-21. After accounting for colony density and location (and/or management strategy), supplemental feeding may improve the 2018-19 model fit (as measured by AIC), but does not improve prediction for 2020-21.

**Table 4.** Multiple regression comparison, with spatial component. Note that management practices are correlated with location. The best model for each statistic is bolded. There is no clear winner amongst the models for all criteria.

| years | stat | Density | Management | Supp. Feed | Dens/Manage | Dens/Supp | Manage/Supp |
| --- | --- | --- | --- | --- | --- | --- | --- |
| 2018-19 | AIC | 1235.52 | 1236.38 | **1214.16** | 1236.05 | **1213.74** | 1218.23 |
| 2018-19 | RelErr | **0.77** | 0.83 | 0.81 | 0.80 | 0.79 | 0.79 |
| 2018-19 | MAE | **3.57** | 3.89 | 3.70 | 3.79 | 3.63 | 3.76 |
| 2018-19 | Rsqr | 0.08 | **0.09** | 0.07 | 0.08 | 0.05 | 0.02 |
| 2020-21 | RelErr | **0.78** | 0.81 | 0.86 | 0.79 | 0.84 | 0.81 |
| 2020-21 | MAE | **3.81** | 3.94 | 4.60 | 3.91 | 4.54 | 4.41 |

**Table 5.** Multiple regression comparison, without spatial component, because management practices are correlated with location. The best model for each statistic is bolded. There is no clear winner amongst the models for all criteria.

| years | stat | Density | Management | Supp. Feed | Dens/Manage | Dens/Supp | Manage/Supp |
| --- | --- | --- | --- | --- | --- | --- | --- |
| 2018-19 | AIC | 1239.06 | 1230.91 | 1226.62 | 1231.11 | 1226.47 | **1221.66** |
| 2018-19 | RelErr | 0.89 | 0.86 | 0.90 | 0.84 | 0.88 | 0.84 |
| 2018-19 | MAE | 3.90 | 3.94 | 3.86 | 3.87 | **3.81** | 3.84 |
| 2018-19 | Rsqr | 0.05 | 0.13 | **0.25** | 0.13 | 0.24 | 0.19 |
| 2020-21 | RelErr | 0.89 | 0.85 | 0.97 | **0.84** | 0.94 | 0.88 |
| 2020-21 | MAE | 4.18 | 4.05 | 4.59 | **4.03** | 4.50 | 4.29 |

Figure 6 and Table 6 show the predicted *Varroa* intensity over the course of a season and the predicted daily growth rate from the multiple regression model which includes Supplemental Feeding (Yes/No), *Varroa* management practice (Monitor + Treat, Treat only, None) and colony density (number of colonies within 5 km).

**Fig 6.**
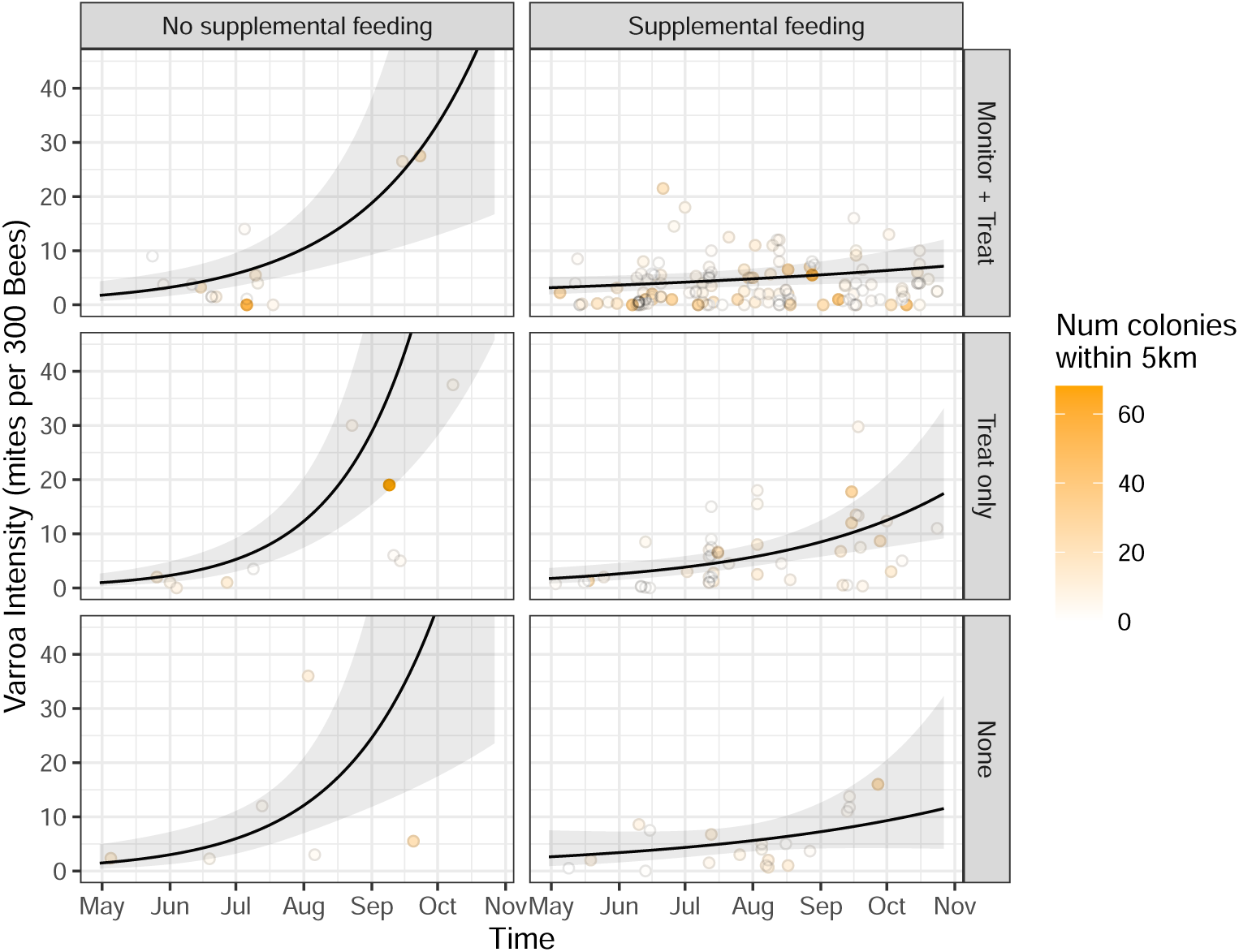
Observed *Varroa* intensity (points) and predicted intensity (line) from multiple regression model without spatial component using 2018-2019 data, by supplemental feeding and management practice categories. Line indicates predicted mean *Varroa* (gray band = 95% confidence interval predicted when number of colonies in 5 km radius is 5 (the median value).

**Table 6.** Predicted *Varroa* intensity and daily *Varroa* growth rate from multiple regression model with spatial component. Values are estimated for a colony density of 5 colonies within 5 km (the median value). May 1 intensity is the mean *Varroa* intensity (mites per 300 bees) predicted for May 1; Growth rate is the daily growth rate of *Varroa* as a multiplicative factor; Growth % is the daily growth rate of *Varroa* as percent change (0-100).

| Management | Feeding | May 1 intensity | Growth rate | Growth % |
| --- | --- | --- | --- | --- |
| Monitor + Treat | No supplemental feeding | 1.76 | 1.019 | 1.93 |
| Monitor + Treat | Supplemental feeding | 3.17 | 1.005 | 0.45 |
| Treat only | No supplemental feeding | 0.96 | 1.028 | 2.78 |
| Treat only | Supplemental feeding | 1.73 | 1.013 | 1.29 |
| None | No supplemental feeding | 1.44 | 1.023 | 2.31 |
| None | Supplemental feeding | 2.60 | 1.008 | 0.83 |

The sample sizes for the “No supplemental feeding” groups are small, leading to large uncertainty bounds; however, the predicted *Varroa* intensity and growth rate are statistically significantly higher (p-value = 0.003) than for the “Supplemental feeding” group. Supplemental feeding seems to have the largest effect size. There is not a significant difference (p-value = 0.484) between the “Monitor + Treat” and “None” management groups, but the “Treat only” group has significantly higher growth rate than “Monitor + Treat” (p-value = 0.031). Table 7 shows the positive relationship between colony density and growth rates in the “Monitor + Treat” with “Supplemental feeding” group. This suggests supplemental feeding along with a *Varroa* monitoring and treatment regime may be the best strategy to keep *Varroa* intensity low throughout the season. However, while the impact of the *Varroa* management strategy appears clear, the impact of supplemental feeding is less clear, particularly as it is less important for predicting 2020-21 outcomes (see also Supplementary Material).

**Table 7.** Predicted *Varroa* intensity and daily *Varroa* growth rate from multiple regression model (with spatial part) assuming “Monitor + Treat” management strategy and “Supplemental feeding.” Values are estimated for varying colony density, from 0 to 60 colonies within a 5 km radius. May 1 intensity is the mean *Varroa* intensity (mites per 300 bees) predicted for May 1; Growth rate is the daily growth rate of *Varroa* as a multiplicative factor; Growth % is the daily growth rate of *Varroa* as percent change (0-100).

| Colony density | May 1 intensity | Growth rate | Growth % |
| --- | --- | --- | --- |
| 0 | 2.14 | 1.011 | 1.13 |
| 1 | 2.05 | 1.012 | 1.16 |
| 2 | 1.97 | 1.012 | 1.19 |
| 5 | 1.73 | 1.013 | 1.29 |
| 10 | 1.40 | 1.015 | 1.46 |
| 20 | 0.91 | 1.018 | 1.79 |
| 60 | 0.16 | 1.031 | 3.12 |

## Discussion

With the goal of understanding the main predictors of *Varroa* infestation in managed apiaries over large spatial areas, we train and test several spatiotemporal models using four years of apiary inspection data in the state of Illinois. Like many ecological datasets, our data have limited resolution in both space and time; this makes validation of mechanistic modeling challenging. Therefore, we use statistical models to probe the risk factors for *Varroa*, including time of year, location, colony density, several environmental factors, and several beekeeper behaviors.

Our baseline model, accounting for only time of year and location shows that *Varroa* intensity grows exponentially in time, as shown in other studies (e.g., [32]), and is also spatially varying. After accounting for location and time of year, nesting quality for wild bees is not predictive of *Varroa* intensity, as expected; our dataset is comprised entirely of managed apiaries. However, it is surprising that other environmental factors such as floral quality and insecticide burden are also not predictive of *Varroa* intensity, as limited environmental nutrition has been shown to exacerbate parasite load [46–49]. That said, the nearly universal practice of supplemental feeding in our dataset may nullify the effects of poor nutrition in the natural environment. Our work aligns with other observational studies that show *Varroa* presence itself erases the influence of any environmental factors [4]; however, in a controlled experiment, it has been found that insecticide exposure increases *Varroa* intensity [50].

After accounting for spatiotemporal effects, increased colony density appears to be a major risk factor for infestation. This observation may be due simply to increased interaction between susceptible and infected bees in the unnaturally high colony densities of modern apiculture [51]. Studies show that bees are more likely to drift in high density apiaries [52] and bees engaged in robbing are more likely to be infected with pathogens [53–57]. In particular, infection with Israeli acute paralysis virus (vectored by *Varroa* mites) has been shown to increase transmission of pathogens by altering infected bee pheromones, allowing bees to slip past susceptible colony defenses [58]. It is also possible that beekeepers themselves may be inadvertently transferring parasites between colonies within their high-density apiaries [59, 60].

After accounting for location and time of year, certain beekeeper behaviors (e.g., supplemental feeding, monitoring and treating for *Varroa*) appear to significantly reduce the *Varroa* growth rate, while treating without monitoring appears to offer no benefit over no treatment. This confirms other studies showing that supplemental nutrition and monitoring reduce parasite load [61]. It is also a logical result; if beekeepers are treating only, they rely on guesswork and luck to treat at the right time for efficacy. That said, some beekeeper behaviors (e.g., monitoring for *Varroa*) are correlated with location. Therefore, it is challenging to know if location itself or beekeeper monitoring is a predictor of infestation.

### Limitations

While ideally a study like this would disambiguate the role of location/time, environmental conditions and beekeeper behaviors in exacerbating or protecting against *Varroa* infestation, sparse and missing data limit our conclusions. Comprehensive, consistent, and time-dense data collected over large spatial areas are not currently available for honey bee colonies. Arguably the most comprehensive data collector is the Bee Informed Partnership (beeinformed.org), but the data are not publicly available, and the organization is reducing its operations.

Another spatial complexity for which we do not account is the common practice of migratory beekeeping, which has also been linked to pollinator disease spread [62–64]. It is speculated that bees shipped thousands of miles to satisfy seasonal pollination demand will experience an increased risk of parasitism and infectious disease compared with colonies that remain in a single location, although rigorous experimental studies have not been performed to verify the observational claim [65]. Pathogen spread through migratory beekeeping remains an understudied phenomenon, despite the practice’s frequency and importance in agriculture [65].

Finally, correlations (even where our evidence is significant) do not imply causation. The existence or direction of causal relationships among time/location, environmental conditions, beekeeper behaviors, and parasite load remain less clear, primarily because causal studies are difficult and expensive to perform across large spatial scales. Both dense ecological data and controlled experiments are needed to resolve these ambiguities. Smaller scale causal studies could identify optimal time(s) to feed and treat in order to minimize disease burden.

## Conclusion

Using a spatiotemporal model of *Varroa* infestation in managed apiaries across the state of Illinois over four years, we test the correlations among parasite intensity, environmental conditions, and beekeeper behaviors. Surprisingly, we find that environmental factors, such as floral quality and insecticide burden, are not predictive of *Varroa* growth. On the other hand, factors largely within beekeeper control (e.g., colony density, supplemental feeding, and parasite monitoring/treatment) predict mite intensity. Smaller colony density and supplemental feeding appear to be protective against *Varroa* growth. Interestingly, while monitoring *and* treating for mites is protective, treating *without* monitoring is no more effective than not treating at all. This is an important result supporting Integrated Pest Management (IPM) approaches.

## Data Availability

All R code and data are publically available at github.com/lfbv/Varroa-Intensity.

## Acknowledgements

The authors thank Marco Ruiz and Meredith Frey for early contributions to modeling. The authors also greatly appreciate Charlotte Blake for scraping Beescape environmental data.

## Supplementary Materials

### Investigating spatial and temporal autocorrelation

After fitting the baseline negative binomial regression model (Equations 2), we examine the residuals for spatial and temporal autocorrelation.

The empirical semivariogram (Figure S1) indicates no detectible spatial autocorrelation even at the smallest lag distance (¡1 km), though it is possible spatial correlation could exist at a finer scale. Additionally, spatial correlation may not be detectable because measurements that were taken close together spatially were not necessarily taken close together in time.

**Fig S1.**
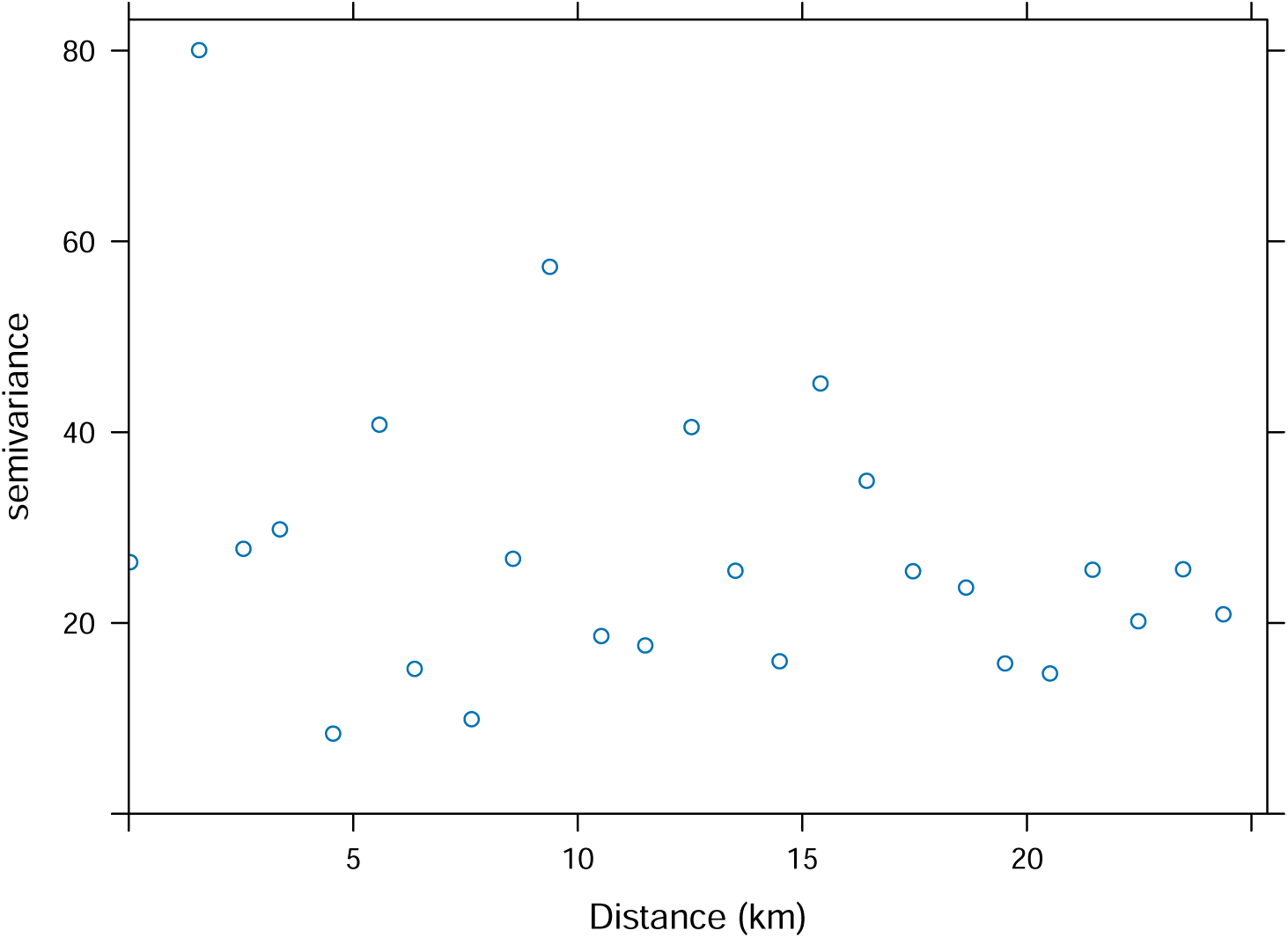
Empirical semivariogram of residuals from baseline model using 2018-19 training data. Bin width is 1 km.

We also examine the temporal autocorrelation for both the residuals and for the measured *Varroa* intensity (Figure S2), which shows no detectable autocorrelation. However, this result only demonstrates that we are unable to detect autocorrelation with the data available, not that no autocorrelation exists. Of the 540 available days of data from the two seasons, *Varroa* intensity is only measured on 134 unique days, and only 61 pairs of consecutive days. Additionally, even if measurements are available on consecutive days, they may be far apart in space, thus making the temporal autocorrelation difficult to detect.

**Fig S2.**
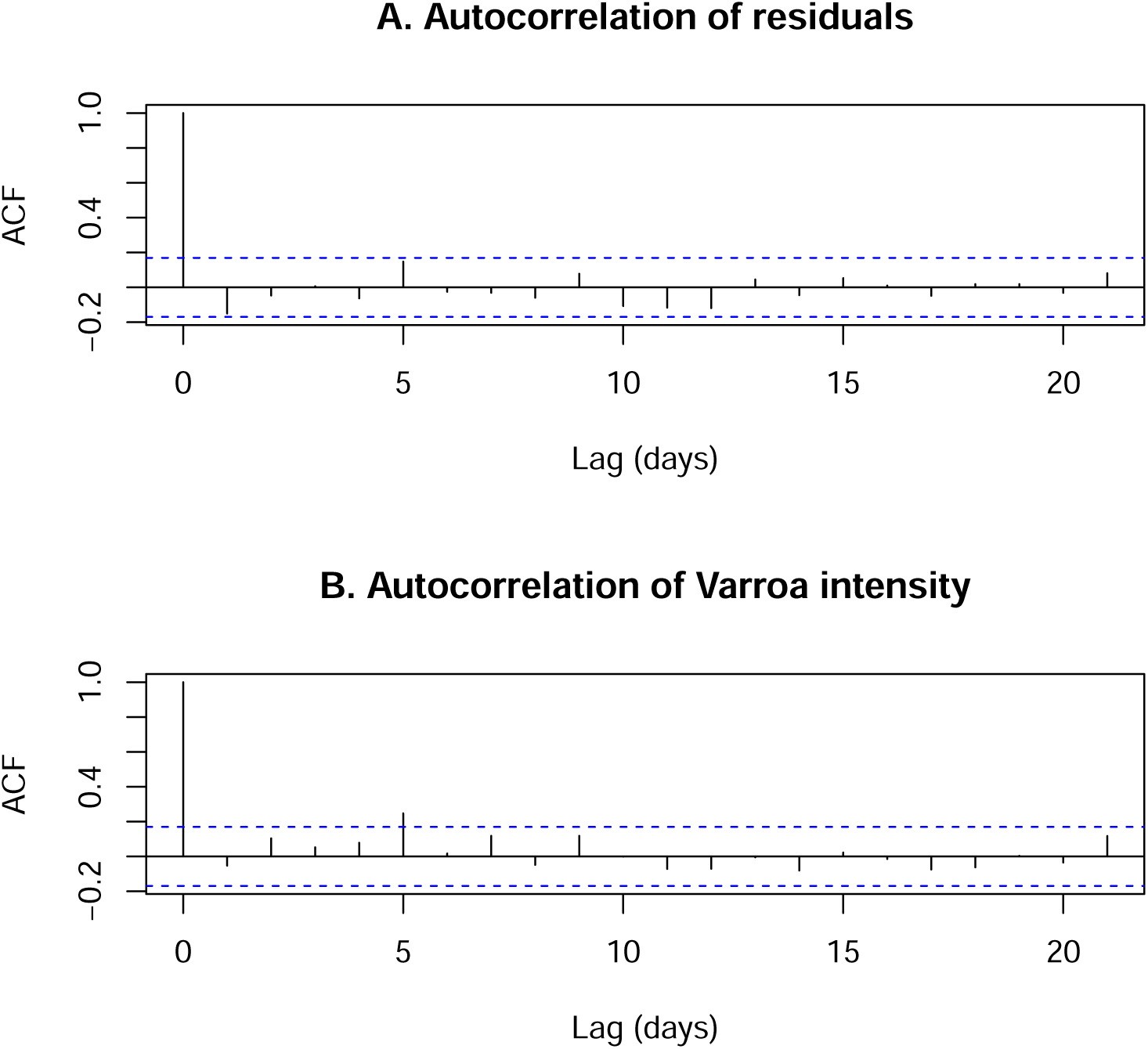
Empirical lagged autocorrelation function (ACF)

### Possible interaction between colony density and time to predict *Varroa* intensity

As described in the section “Impact of environmental factors”, colony density is not statistically significant in a single variable additive model, but the interaction between colony density and time of year does appear significant. Figures S3 and S4 show that this association may be driven by high late-season *Varroa* intensity for apiaries with very high nearby colony density. Predicted intensity is similarly low for all values of colony density until mid-August, when the predicted intensity for the higher density regions increases rapidly.

**Fig S3.**
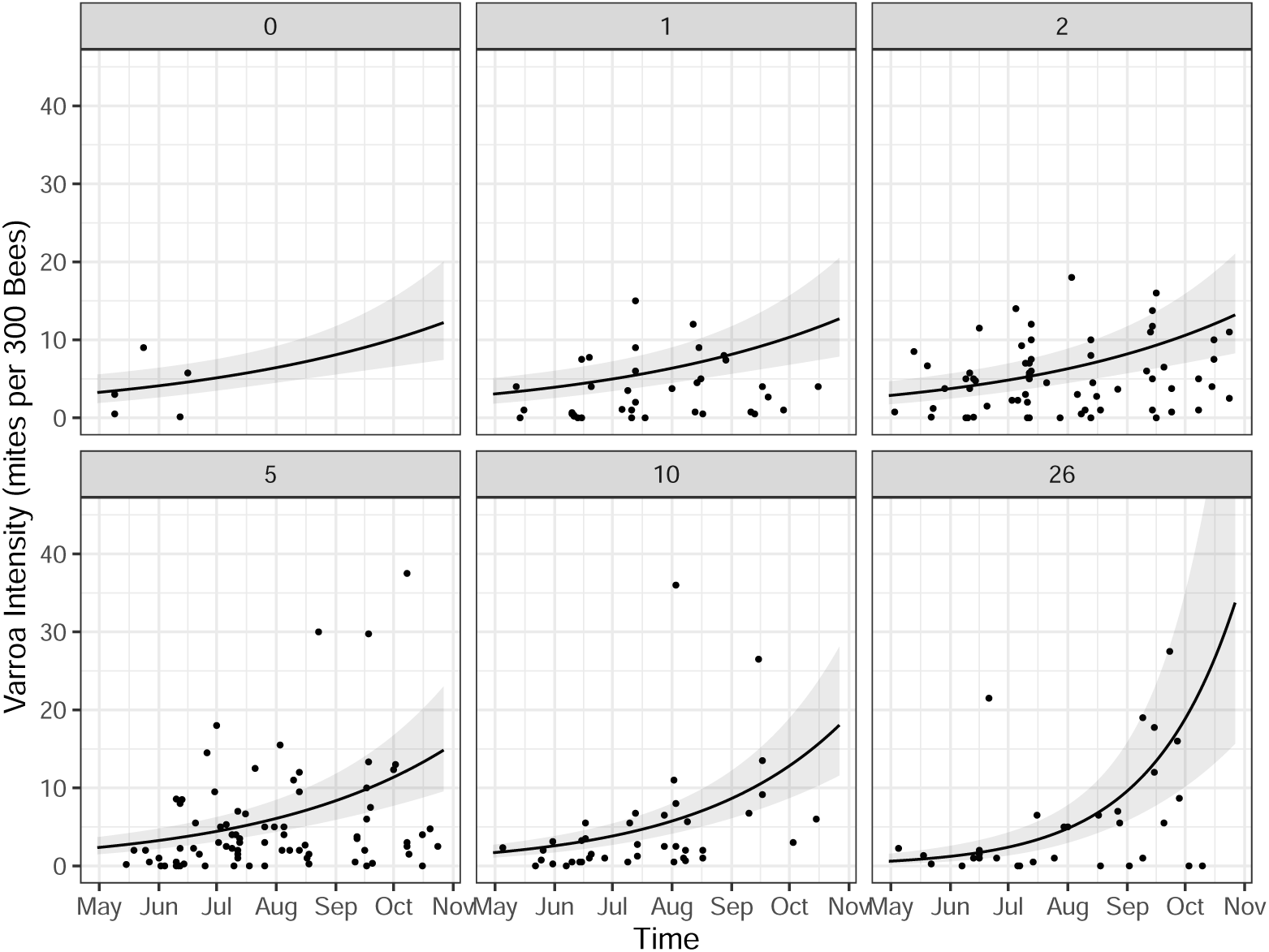
Varroa intensity over time for different levels of colony density. Each point represents one apiary, panel labels indicate approximate number of colonies within a 5km radius. Black line indicates the *Varroa* intensity in mites per 300 bees as predicted from a model with interaction between day of year and colony density, along with gray 95% confidence band.

**Fig S4.**
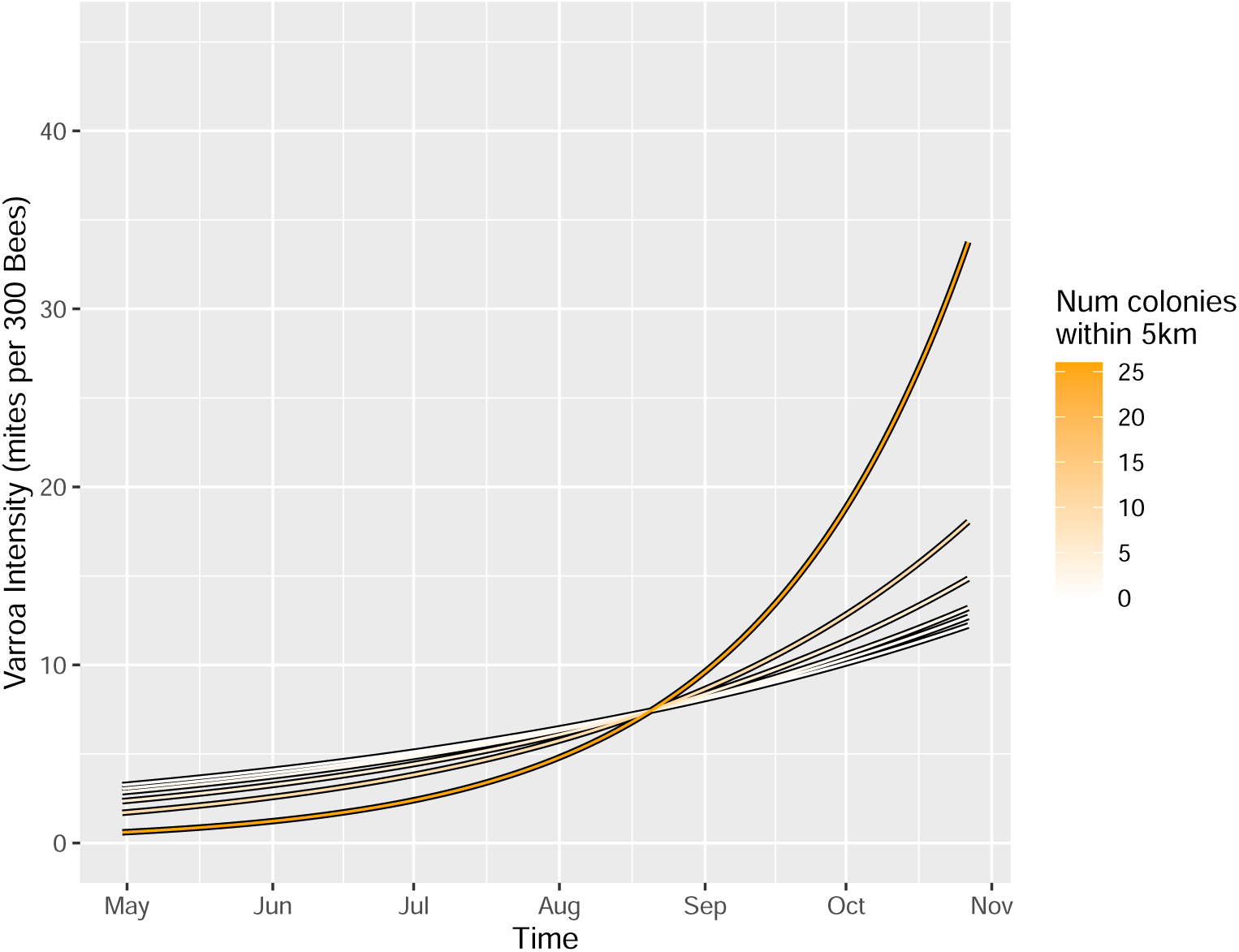
Predicted *Varroa* intensity from model with interaction of colony density and day of year. These are the same models shown in Figure S3.

